# A study of hyperelastic continuum models for isotropic athermal fibrous networks

**DOI:** 10.1101/2022.06.28.497976

**Authors:** Dawei Song, Assad A Oberai, Paul A Janmey

## Abstract

Many biological materials contain fibrous protein networks as their main structural components. Understanding the mechanical properties of such networks is important for creating biomimicking materials for cell and tissue engineering, and for developing novel tools for detecting and diagnosing disease. In this work, we develop continuum models for isotropic, athermal fibrous networks by combining a single-fiber model that describes the axial response of individual fibers, with network models that assemble individual fiber properties into overall network behavior. In particular, we consider four different network models, including the affine, three-chain, eight-chain, and micro-sphere models, which employ different assumptions about network structure and kinematics. We systematically investigate the ability of these models to describe the mechanical response of athermal collagen and fibrin networks by comparing model predictions with experimental data. We test how each model captures network behavior under three different loading conditions: uniaxial tension, simple shear, and combined tension and shear. We find that the affine and three-chain models can accurately describe both the axial and shear behavior, whereas the eight-chain and micro-sphere models fail to capture the shear response, leading to an unphysical zero shear moduli at infinitesimal strains. Our study is the first to systematically investigate the applicability of popular network models for describing the macroscopic behavior of athermal fibrous networks, offering insights for selecting efficient models that can be used for large-scale, finite-element simulations of athermal networks.

## 1. Introduction

Many biological materials consist of fibrous protein networks as their main structural components, and such networks play a critical role in determining their mechanical properties and other functions [1, 2, 3, 4, 5, 6]. For instance, the extracellular matrix (ECM) of tissues contains a random network of collagen and elastin [7, 8]. Changes in the structure of these networks can give rise to abnormal mechanical properties of tissues that disrupt cell mechanosensitive responses and, in turn, initiate or exacerbate pathologies [9, 10]. Blood clots are composed of networks of fibrin, whose structure and mechanical properties are crucial for preventing bleeding [11, 12, 13, 14]. Thus, understanding and quantifying the mechanical behavior of fibrous networks is important not only for designing biomimetic materials that serve as physiologically relevant environment for cell engineering, but also for developing novel diagnostic techniques for disease [15, 16]. With this as motivation, this paper is concerned with the development of continuum models for fibrous networks.

Fibrous networks can be classified into thermal and athermal networks, depending on the nature of the constituent fibers [3]. A network is said to be thermal when its fibers are subjected to significant thermal fluctuations, such that the fiber’s persistence length (i.e., the length over which the fiber appears straight in the presence of Brownian forces) is much smaller or comparable to its contour length [2]. Examples include molecularly crosslinked, flexible networks like rubbers and polyacrylamide gels, in which the fiber persistence length is far less than its contour length and the fiber appears as a random coil [17]. Another example is given by semiflexible, cytoskeletal networks like actin and intermediate filament networks, in which the fiber has a persistence length comparable to its contour length or the distance between network crosslinks [2]. In this case, the bending energy of the fiber just outcompetes the entropic tendency of the fiber to crumple into a random coil, and the fiber exhibits small but significant thermal fluctuations around a straight conformation. On the other hand, a network is said to be athermal when its fibers are large enough or rigid enough to be negligibly affected by thermal fluctuations, with the fiber’s persistence length being much larger than its contour length [3, 18]. Examples of athermal networks include collagen networks in the ECMs and fibrin networks in blood clots. These athermal fibers often behave mechanically like slender elastic beams.

Continuum modeling of fibrous networks treats the network as a mechanical continuum, with the goal of developing a constitutive relation that describes the macroscopic response of the network in relation to the properties of the constituent fibers and the network structure [19]. Such modeling efforts typically comprise two steps. The first step is to derive a force-extension relation for single fibers along their axial direction (i.e., the direction connecting the two ends of the fiber). For flexible fibers, entropic models of Gaussian or Langevin type are usually used [17]. The Gaussian model assumes that the fiber is a freely jointed chain of rigid rods, being infinitely stretchable without approaching its fully extended state; this results in a linear force-extension relation. In contrast, the more realistic Langevin model accounts for the finite stretchability of the fibers yielding a strain-stiffening force-extension relation. For semiflexible fibers, different models have been proposed (e.g., [20, 21]) building upon the classical worm-like chain model [22], treating fibers as continuous and smooth filaments with well-defined local curvatures. Then, fiber behavior can be described by the free energy of bending of fibers induced by thermal fluctuations. As a fiber is stretched, the amplitude of its transverse thermal undulations decreases, causing a decrease in available fiber conformations and thus an entropic strain-stiffening of the fiber. Different from the above entropic models, models for athermal fibers are intrinsically energetic, so that the force-extension relation can be derived from the stored elastic energy of the fibers. These fibers often exhibit a linear elastic or strain-stiffening response under axial tension, but can buckle when subjected to axial compression, thus having a limited compression resistance [3, 23].

The second step of continuum modeling is to develop a network model that relates the stretch of individual fibers to the macroscopic network deformation, so that the overall response of the network can be computed by adding contributions from fibers over all orientations. Over the years, several network models have been developed for isotropic fibrous networks. These network models can be categorized into affine and non-affine models. The affine model (also called the full network model) [17, 24, 25] considers fibers isotropically distributed in the orientation space and assumes that the fibers deform according to the macroscopic deformationof the network (Fig. 1(a)), i.e., the axial stretch of any fiber is taken to be identical to the corresponding macro-stretch of the continuum along the fiber’s direction.

**Figure 1:**
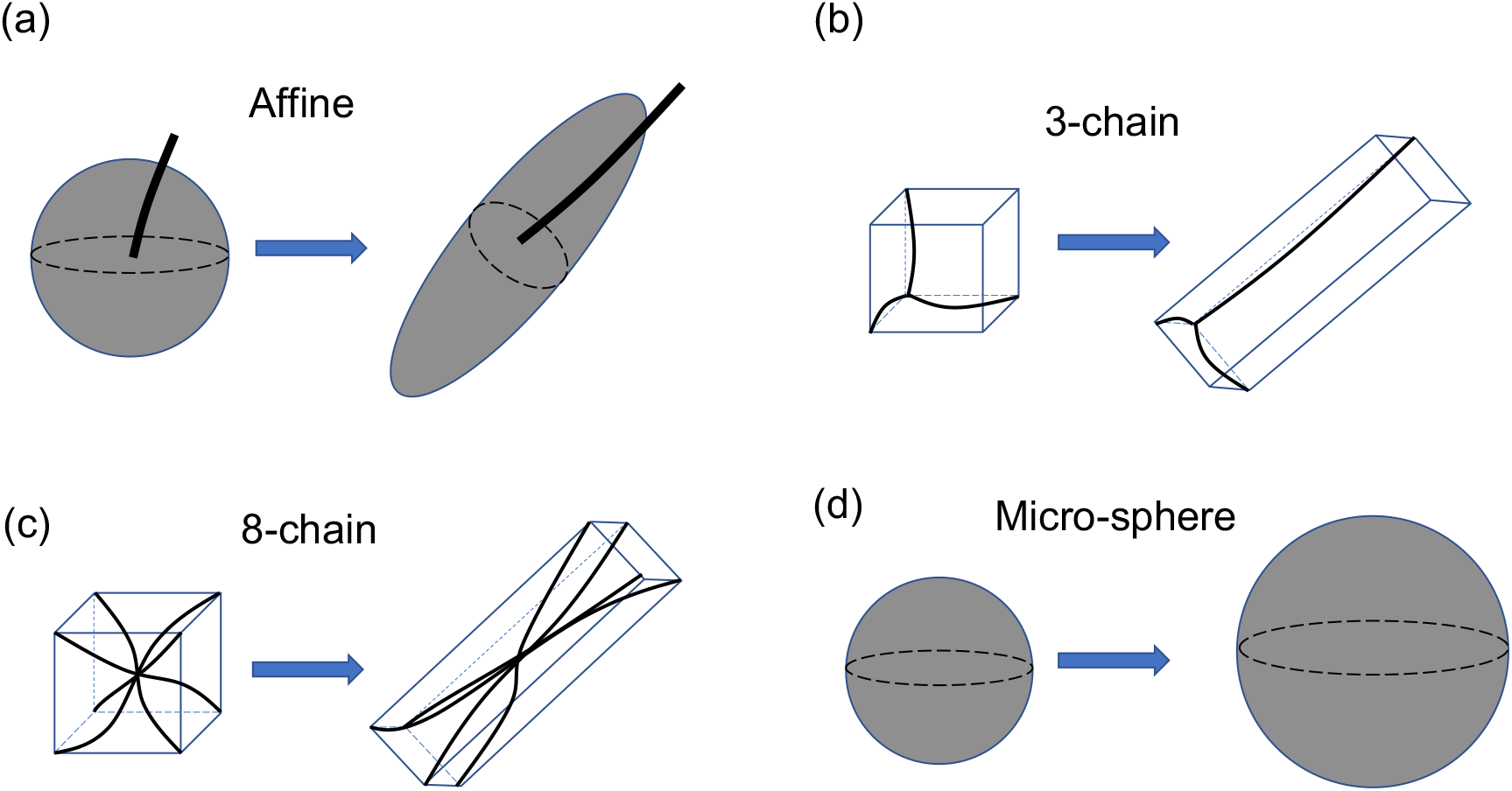
Schematics for the geometry and deformation of different network models under simple shear. (a) Affine model. Each point on the spherical surface represents a unique fiber orientation, with only one representative fiber shown in the figure. (b) Three-chain model, where each fiber is aligned with one of the edges of the unit cell. (c) Eight-chain model, where each fiber connects the center of the unit cell to one of the cell vertices. (d) Micro-sphere model. Each point on the spherical surface represents a unique fiber orientation, with all the fibers stretched to the same degree. The rotation of the fibers, however, is not well defined; thus, fiber deformation is not shown in the figure.

The development of non-affine models, however, is motivated by the fact that the true fiber deformation is usually non-affine [2, 3]. One type of such non-affine models is the unit-cell model [19], which idealizes the actual structure of the network as a periodic array of unit cells, each of them containing a certain number of fibers with prescribed arrangements. The most popular unit-cell models are perhaps the three-chain model [26] and the eight-chain model [27]. In the three-chain model, the unit cell is initially cubic containing three mutually orthogonal fibers along its edges (Fig. 1(b)). Under a prescribed macroscopic deformation, the three fibers are taken to be aligned with the principal directions of the deformation, and the stretch of the fiber is determined by the corresponding principal stretch value. The eight-chain model has a setting similar to that of the three-chain model, the key difference being that the unit cell contains eight fibers connecting the center of the cell to the eight cell vertices (Fig. 1(c)). Due to the symmetry of such a network structure, the stretch of all the eight fibers equals the root mean square of the principle stretch values. Another type of non-affine model is the so-called micro-sphere model [28], which also considers fibers isotropically distributed in the orientation space (Fig. 1(d)). In this model, however, the axial stretch of a given fiber differs from the corresponding macro-stretch of the continuum, and is obtained by minimizing the average free energy of the fibrous networks. This minimization procedure yields an effective axial stretch of all the fibers to be the *p*-root average of the macro-stretch over all orientations, where *p* is a material parameter describing the network’s non-affinity. For the special case of *p* = 2, the micro-sphere model recovers the eight-chain model.

These different network models, along with appropriately chosen force-extension relations for individual fibers (e.g, Langevin model for flexible fibers, and worm-like chain model for semi-flexible fibers), have been extensively used to study the overall behavior of thermal networks, such as rubber [19, 25], actin [20, 29, 30], and intermediate filament networks [31]. These networks are either incompressible themselves (e.g., dry rubber networks), or embedded in an incompressible fluid (e.g., actin networks in cytosol). Since these thermal networks typically have small pore sizes, fluids do not have enough time to migrate through the network on a time scale of seconds to minutes. Therefore, in these scenarios, thermal networks can be modeled as *incompressible*, hyperelastic materials. In particular, for the various network models described above, fairly good agreement between model predictions and experimental data was generally found. Compared with the affine and three-chain models, the eight-chain model was found to be better at simultaneously capturing the behavior of rubber networks under different loading conditions (e.g., uniaxial tension, biaxial tension, and simple shear) [27, 19]. The micro-sphere model, which includes the eight-chain model as a special case, further improves model capacity and provides better predictions for the behavior of rubber and actin networks [28].

Despite the extensive and systematic study of network models for thermal networks, it remains unclear if these models can be extended to accurately describe the macroscopic behavior of *athermal* networks, such as collagen and fibrin networks.Different from thermal networks, athermal networks often have a much more open structure with large pore sizes; consequently, fluids can flow in and out of the networks on a similar time scale, causing significant changes in the network volume [18, 32]. However, in this work we do not consider the dynamic network response induced by fluid flow; instead, we are interested in the quasi-static behavior of the networks after fluid flow has completed and the network has reached an equilibrium state, in which case the athermal networks can be modeled as *compressible*, hyperelastic materials. In particular, we ask the following question: given that the affine, three-chain, eight-chain, and micro-sphere models have been widely used to describe the mechanical properties of incompressible, thermal networks, can they be further generalized to capture the quasi-static behavior of compressible, athermal networks?

To answer this question, we develop various continuum models for athermal networks by combining each of the above network models (the affine, three-chain, eight-chain, and micro-sphere models) with a single-fiber model that has been found to successfully capture the axial response of athermal fibers. This single-fiber model was first developed by Steinwachs et al. [5], assuming that the fiber can stiffen under axial tension and soften under axial compression. Further, to account for the compressibility of the network, we incorporate into our models a volumetric energy term that can describe nonlinear volumetric responses of the network, following the earlier work of [33]. Thereafter, we systematically evaluate the models’ performance by assessing how well they fit available experimental data for collagen and fibrin networks subjected to various loading conditions, including uniaxial tension, simple shear, as well as combined tension and shear. We also analyze why different models may have different capacities in capturing the true behavior of athermal networks. Finally, we end with the conclusions of this study.

## 2. Continuum models for athermal fibrous networks

In this section, we develop various continuum models for isotropic, athermal fibrous networks. As already discussed in the Introduction, these models contain two parts: (i) a single-fiber model that describes the axial force-extension response of individual fibers, and (ii) a network model that describes the structure and kinematics of the networks, as well as determines the overall network behavior by adding contributions from all the fibers. Next, we discuss these two components.

### 2.1. Single-fiber models

We characterize the axial response of individual fibers using the model developed by Steinwachs et al [5]. This model has been used to successfully describe the behavior of athermal collagen fibers, capturing the asymmetric response of fibers to tension and compression. In particular, the fiber is assumed to be wavy in its unloaded state and to exhibit three distinct regimes in response to external forces. When the fiber is axially stretched, it has an initially constant stiffness, up to the point where the fiber is fully straightened. Upon further stretching, the fiber gradually stiffens with a stiffness that increases with tensile strain. On the contrary, when the fiber is axially compressed, it softens with a stiffness that decreases with compressive strain. More specifically, the fiber’s differential stiffness, *κ*_*f*_, is given by

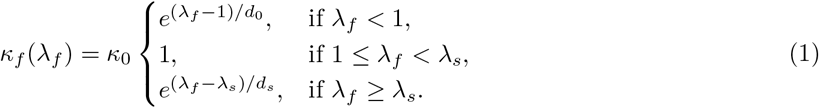

In the above expression, *λ*_*f*_ is a kinematic variable denoting the axial stretch of the fiber, as given by the ratio between the end-to-end length of the deformed fiber and that of the undeformed fiber. Moreover, *κ*_0_ is the initial stiffness of the fiber, *λ*_*s*_ is the critical stretch at the onset of fiber stiffening (i.e., when the fiber is just straightened), *d*_*s*_ is a coefficient describing the exponential stiffening of the fiber, and *d*_0_ is a coefficient describing the exponential softening of the fiber; these material parameters define the material properties of a single fiber and are determined from experimental data. We note that *d*_*s*_ (or *d*_0_) can be interpreted as the strain scale of fiber stiffening (or softening), where smaller values of *d*_*s*_ (or *d*_0_) correspond to faster rate of stiffening (or softening).

For later use, we also determine the axial force, *F*_*f*_, and the strain energy, *w*_*f*_, associated with a single fiber. Following earlier studies, we assume that the fiber is force-free and has a zero strain energy in its unloaded state. Then, making use of the relation 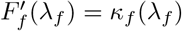 and 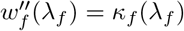, we may compute *F*_*f*_ and *w*_*f*_ by integrating *κ*_*f*_ with respect to *λ*_*f*_ once and twice, respectively. Consequently, we have the results that

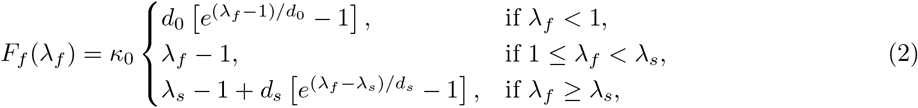

and

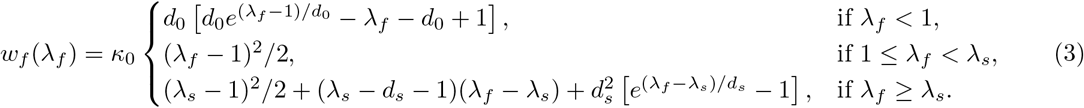

We note that several other models, such as the linear model of [34, 35] and the exponential model of [36, 37], have also been used to describe the axial response of individual fibers. Those models typically assume that fibers buckle immediately under compression and cannot sustain compressive forces, as would be the case if *d*_0_ → 0 in (1)-(3). However, we do not make such an assumption; instead, we treat *d*_0_ as a material parameter and determine it by fitting the model to experimental data. As will be seen below, *d*_0_ determined from experimental data is indeed very close to 0.

### 2.2. Network models

In this subsection, we present network models for isotropic fibrous networks. We also combine each of these network models with the above single-fiber model to develop full continuum models for athermal fibrous networks.

As already mentioned in the Introduction, we focus on the quasi-static behavior of athermal networks, which can be modeled as *compressible*, hyperelastic solids. Then, the mechanical behavior of the networks can be fully described by a strain energy density function, *W* [38]. Following earlier works on compressible hyperelastic solids [19], *W* may be written in an additive form:

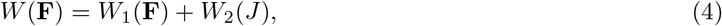

where **F** = ∂**x**/∂**X** is the deformation gradient tensor, with **x** and **X** being the positions of a material point in the deformed and undeformed configurations, respectively, *J* = det(**F**) is the Jacobian, *W*_1_ is the strain energy density associated with a chosen network model (see below), and *W*_2_ is the volumetric strain energy density due to volume changes. Different models for *W*_2_ have been proposed in the literature; interested readers are referred to the excellent book of Anand and Govindjee [39] for a comprehensive review. Here we employ the model developed by Bischoff et al. [33], which has the capacity to describe general nonlinear volumetric response of a material. In this model, *W*_2_ is given by

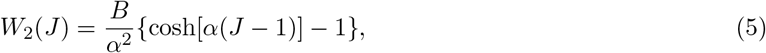

where *B* determines the bulk modulus of the material, and *α* describes the nonlinearity of the volumetric response. Note that *W*_2_ is empirical in nature, and both *B* and *α* are material parameters to be determined from experimental data.

*W*_1_ in (4) denotes the strain energy density predicted by a chosen network model. Such models account for the network structure, first computing the stretch of individual fibers from the macroscopic network deformation, and then computing the total strain energy of the network by summing up the energy of all the fibers. In the following, we will briefly summarize various network models, including the affine, three-chain, eight-chain, and micro-sphere models. As will be seen, these models differ in how they describe the isotropic network structure, as well as in how they compute the axial stretch of individual fibers from the macroscopic network deformation.

Before introducing the network models, we note that we will make the following assumptions that will apply to all models. First, although different fibers in a random network generally have different shapes (e.g., the end-to-end length and the contour length), we assume, for simplicity, that all the fibers have the same shape. This shape can be interpreted as the *average* shape of all the fibers, as can be described by the average end-to-end length and the average contour length of all the fibers. Second, in order to ensure that the fibrous networks are stress-free in the undeformed configuration, we assume that all the fibers are initially force free. In reality, the initial state of a fiber (i.e., force free, pre-streched, or pre-compressed) depends sensitively on the local polymerization conditions of the network. A detailed account of these effects, however, is beyond the scope of this work.

#### 2.2.1. Affine model

The affine model accounts for all the fibers isotropically distributed in space, and assumes that these fibers deform affinely with the continuum [17, 19] (Fig. 1(a)). As a result, the axial stretch of each fiber, 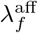, is given by the macro-stretch of the continuum along the direction of that fiber, i.e.,

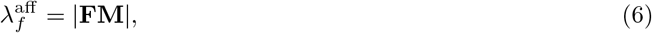

with **M** being the unit vector along the fiber orientation in the undeformed configuration, and |·|denoting the norm of a given quantity. Thus, the strain energy density associated with the affine model can be computed by integrating the strain energy of fibers over the entire orientation space:

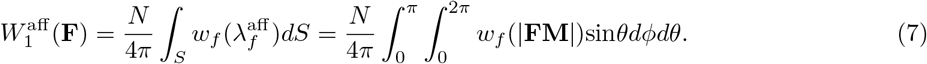

Here *N* is the number of fibers per unit volume of the undeformed network, *S* is the surface of a unit sphere representing the orientation space, with each point on *S* denoting a unique fiber orientation, and *θ* and *ϕ* are, respectively, the polar angle and the azimuthal angle in a spherical coordinate. Note that **M** can be expressed in terms of *θ* and *ϕ* as *M* = [sin*θ*cos*ϕ*, sin*θ*sin*ϕ*, cos*θ*]^*T*^.

#### 2.2.2. Three-chain model

In the three-chain model [3, 26], the network is idealized as a periodic array of initially cubic unit cells, each containing three fibers located along the edges of the cell (Fig. 1(b)). These fibers deform with the unit cell, such that the end-to-end vector of a fiber always coincides with the corresponding cell edge. For a prescribed network deformation, the unit cell deforms into a general cuboid, with the cell edges (or the fibers) always aligned with the principal directions of the deformation, and the stretches of the edges (or the fibers) equal to the corresponding principal stretch values. As such, the strain energy density associated with the three-chain model can be computed by summing up the strain energy of the three classes of fibers:

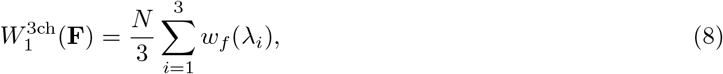

where the *λ*_*i*_ (*i* = 1, 2, 3) denote the principal stretches associated with the deformation gradient **F**.

The deformation of fibers predicted by the three-chain model is generally non-affine. This can be easily seen when the network is subjected to simple shear deformation. As the shear deformation progresses, the three classes of fibers rotate simultaneously while remaining mutually orthogonal (Fig. 1(b)) This is in contrast to what would happen if these fibers deformed affinely with the continuum, in which case the angles between these fibers would change during the course of the simple shear deformation. Finally, we note that the three-chain model can be viewed as a method of *sampling* the rotation and stretching of fibers along the three principal directions of the deformation.

#### 2.2.3. Eight-chain model

The eight-chain model is similar to the three-chain model, the main difference being that there are eight fibers connecting the center of the unit cell to each of the eight cell vertices [27] (Fig. 1(c)). Due to the symmetry of this network structure, the end-to-end lengths of all the fibers equal half of the length of the unit cell’s body diagonal. Thus, the axial stretch of each fiber is given by the root mean square of the principal stretch values, i.e.,

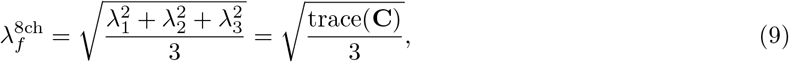

where **C** = **FF**^T^ is the right Cauchy-Green deformation tensor. Consequently, the strain energy density associated with the eight-chain model is given by

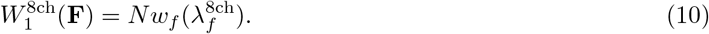

Fiber deformation predicted by the eight-chain model is again non-affine, as can be also seen when the network undergoes simple shear deformation. If the deformation of fibers were affine, then as shear progresses some of the fibers would be stretched whereas the rest would be compressed. This scenario, however, differs from the predictions of the eight-chain model, where all the fibers are stretched to the same degree (see (9) and Fig. 1(c)), implying that fiber deformation is generally non-affine for this model. We also note that the eight-chain model can be viewed as a method of sampling the rotation and stretch of fibers in eight directions relative to the principal directions of the deformation.

#### 2.2.4. Micro-sphere model

The original micro-sphere model [28] considers all the fibers isotropically distributed in space (Fig. 1(d)), with each fiber subjected to a tube constraint. This tube constraint accounts for the conformational constraints placed on a fiber by its neighbors, having important ramifications on the fiber conformational entropy. However, since in this work we focus on athermal networks, whose mechanical response is dominated by the fiber elastic energy and is negligibly affected by its conformational entropy, we expect the tube constraint to have no significant effects on the mechanical behavior of the network. Thus, we neglect the tube constraint. Similar simplifications have also been used in [30], where the authors found that typical parameters for the tube constraint have a minor influence (< 4%) on the final result at small to intermediate strains (< 20%).

The micro-sphere model incorporates non-affine fiber deformations by allowing axial fiber stretches to fluctuate around the corresponding macro-stretches of the continuum. In particular, the axial stretches of fibers are determined by minimizing the network strain energy density, under the *ad hoc* constraint that the *p*-root average of axial fiber stretch equals that of the macro-stretch. Here *p* is a material parameter describing the non-affinity of the network. This minimization procedure leads to the result that the axial stretches of all the fibers equal the *p*-root average of the macro-stretch, i.e.,

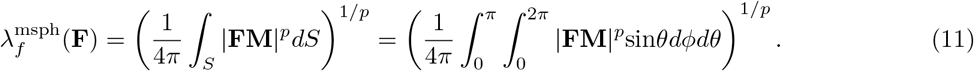

Here we recall that *S* denotes the surface of a unit sphere, where each point on *S* represents a unique fiber orientation, **M**. Thus, the strain energy density associated with the micro-sphere model is given by

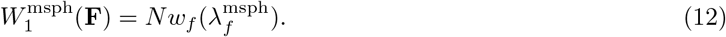

Note that when *p* = 2, equations (11) and (12) can be shown to reduce to (9) and (10), respectively, so that the micro-sphere model recovers the eight-chain model.

In summary, the hyperelastic response of the compressible athermal networks can be characterized by the strain energy density *W* in (4), with *W*_2_ given by (5), and *W*_1_ given by (7), (8), (10), or (12), depending on the choice of the network model. Then, the first Piola-Kirchhoff stress, **P**, of the fibrous network is given by

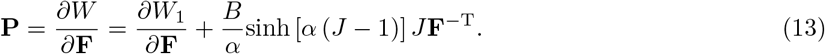

##### Remarks

1. In literature, the names “affine,” “three-chain,” “eight-chain,” and “micro-sphere” have been used to describe different types of network models, which are independent of the choice of single-fiber models. However, since we have combined these network models with the single-fiber model in Section 2.1 to develop full continuum models, we will henceforth use these names to describe the corresponding continuum models.
2. Both the affine and micro-sphere continuum models require the computation of two-dimensional integrals over the surface of a unit sphere (see, e.g., (7) and (11)). These surface integrals are computed by discretizing the surface with 808 triangular elements and employing linear finite element interpolations. This number was found to be sufficient to guarantee convergence [40, 41].
3. The affine, three-chain, and eight-chain continuum models each contain a set of six material parameters, **c** = {*K, d*_0_, *d*_*s*_, *λ*_*s*_, *B, α*}, where *K* = *Nκ* has been treated as a single material parameter (recall that *κ* is the stiffness of individual fibers at small strains, and *N* is the number density of fibers). On the other hand, the micro-sphere continuum model contains a set of seven material parameters, **c** = *{K, d*_0_, *d*_*s*_, *λ*_*s*_, *B, α, p}*.

## 3. Results and Discussion

In this section, we evaluate the performance of the affine, three-chain, eight-chain, and micro-sphere models in describing the mechanical properties of athermal networks, including collagen and fibrin networks. Specifically, we test how well these models capture the network behavior under three different loading conditions, including (i) uniaxial tension, (ii) simple shear, and (iii) combined tension and shear. These loading conditions are commonly used to measure the mechanical properties of fibrous networks. For each loading condition, we fit each of our models to available experimental data (for both collagen and fibrin networks), and assess the accuracy of the model by quantifying the mismatch between model predictions and experimental measurements.

### 3.1. Uniaxial tension

In this subsection, we examine the ability of different models to capture the response of fibrous networks under uniaxial tension. In particular, we compare model predictions with the experimental data of Roeder et al. [42] for collagen networks, and with that of Purohit et al. [43] for fibrin networks.

In standard uniaxial-tension tests, network samples are stretched along a given direction and are allowed to deform freely in directions perpendicular to the loading direction. Thus, we assume that the networks undergo a homogeneous deformation, with the deformation gradient given by

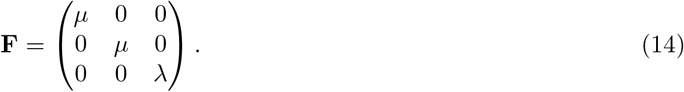

Here *λ* denotes the axial stretch along the loading direction, and *µ* denotes the corresponding lateral stretch, which is chosen in such a way that the lateral stresses predicted by a given model are zero. (Without loss of generality, we have assumed that loading is applied in the *z* direction.) Then, for each model we may compute the axial stress, *P*_*zz*_, as a function of the axial stretch, *λ*, using (13).

We fit the above predicted stress-stretch (*P*_*zz*_-*λ*) response of the network to available experimental data by solving a minimization problem, with the goal of finding the set of material parameters, **c**, that minimizes the cost function

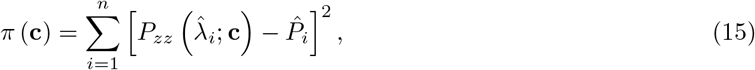

where 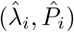 (*i* = 1 *n*) are the pairs of experimental data for the axial stretch and axial stress, respectively, with *n* denoting the total number of experimental data points. We note that under uniaxial tension, most fibers in a network are extended; thus, network behavior is governed mostly by fiber tensile properties, and not by their compressive properties. Consequently, uniaxial-tension data alone is insufficient for determining material parameters that describe compressive responses, such as *d*_0_ in equation (1). Thus, in the above minimization problem we set *d*_0_ = 1 *×* 10^−2^ for all models, and solve for the rest of the material parameters. This choice of *d*_0_ is motivated by the fact that athermal fibers typically buckle easily under compression, so that *d*_0_ (which describes the strain scale of softening) is expected to be very close to 0.

We evaluate the ability of each model to capture the true network response by computing the relative error, which describes how closely the predicted network response matches the measured network response. In particular, we define the relative error as

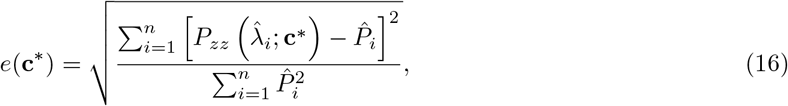

where **c**\* is the set of optimal material parameters that minimizes the cost function (15).

Figure 2 presents the stress-stretch curves predicted by different models when fitted to the experimental data for collagen and fibrin networks. The optimal material parameters obtained from data fitting, as well as the relative errors, are shown in Tabs. S1 and S2 in the Supporting Materials. We observe that for both the collagen and fibrin networks, the slope of the stress-stretch curve (i.e., the tangent modulus of the material) increases with stretch, indicating a nonlinear strain-stiffening response of these networks. This is because as deformation progresses, fibers rotate towards the direction of the maximum principal stretch (i.e., the uniaxial loading direction), leading to realignment along that direction, and thus to strain stiffening of the network. This mechanism of fiber realignment is purely geometric, independent of the material properties of individual fibers. If individual fibers also stiffen with strain, this effect would further enhance the strain-stiffening response of the network. Further, we find that the nonlinear stress-strain curve can be captured fairly well by all the continuum models, with relative errors below 14% (see the Supporting Materials).

**Figure 2:**
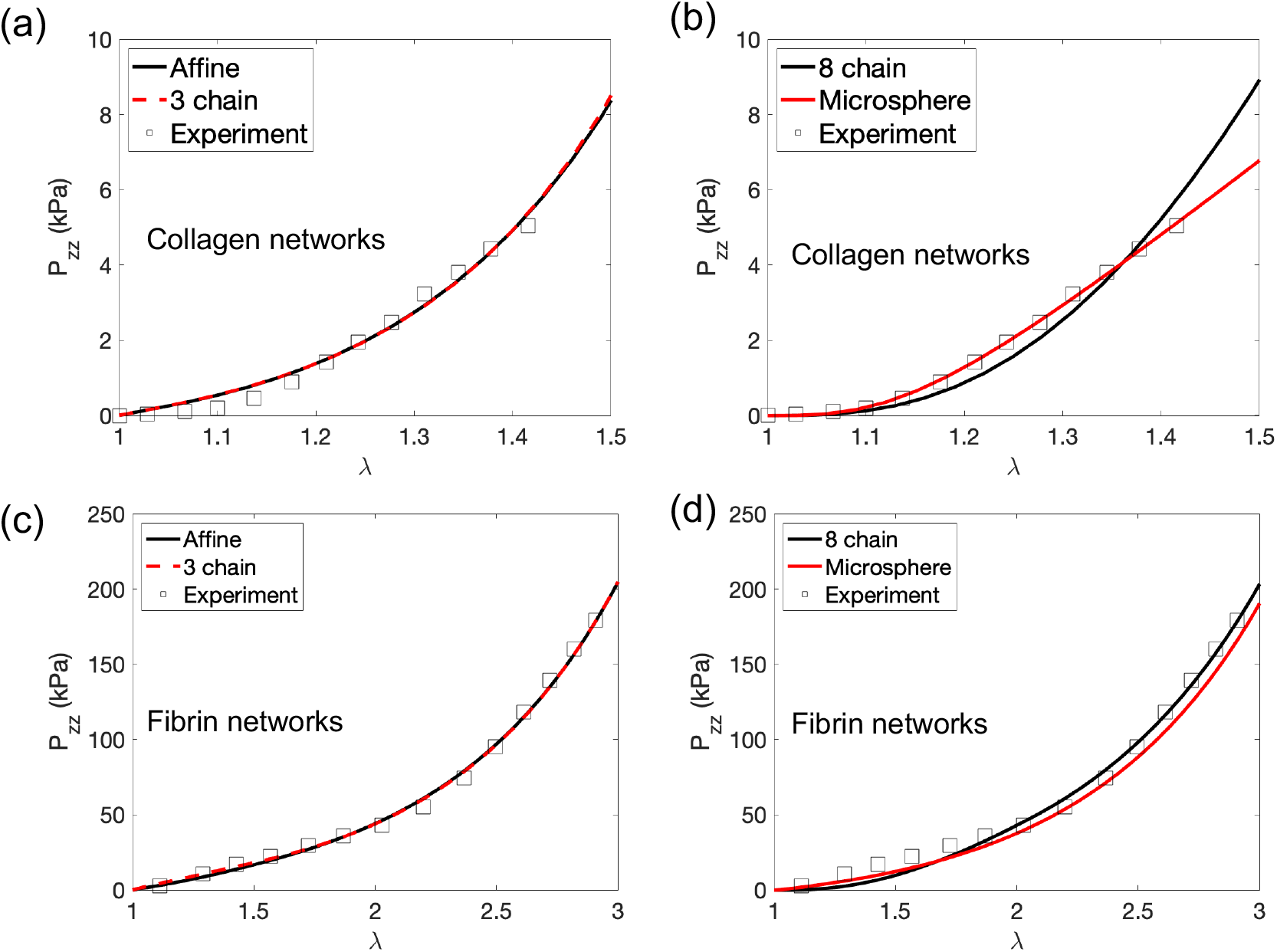
Axial stress *P*_*zz*_ as a function of axial stretch *λ*. Results predicted by the (a) affine and 3-chain models, as well as by the (b) 8-chain and micro-sphere models, are shown for collagen networks. The experimental data of Roeder et al. [42] is also displayed. The corresponding results are also shown for fibrin networks, including predictions of the (c) affine and 3-chain models, as well as of the (d) 8-chain and micro-sphere models. The corresponding experimental data of Purohit et al. [43] is also shown.

### 3.2. Simple shear

In this subsection we test the ability of various models to describe the network behavior under simple shear. To do this, we compare model predictions with the experimental data of Storm et al. [31] for collagen networks and that of van Oosten et al. for fibrin networks [32].

The deformation gradient for simple shear can be written as

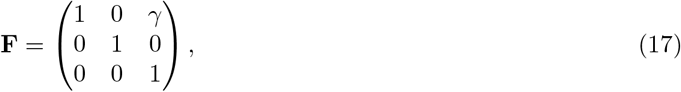

where *γ* denotes the amount of shear. (Here we have assumed, without loss of generality, that shear is applied in the *x*-*z* plane along the *x* direction.) Thus, for each model we can compute the shear stress, *P*_*xz*_, together with the nominal shear modulus, *G* = *P*_*xz*_/*γ*, as a function of *γ*, using (13). We fit the predicted modulus-strain (*G*-*γ*) curve to the available experimental data (for both collagen and fibrin networks) by solving a minimization problem analogous to the problem in Section 3.1. We also evaluate the relative errors using an expression that is similar to (16).

Since simple shear deformation is volume conserving (*J* = 1), the last term in (13) vanishes, implying that network behavior under simple shear is insensitive to the values of *B* and *α* (which describe how a network responds to volume changes). As a consequence, the values of *B* and *α* cannot be inferred from shear response. Motivated by this observation, when performing curve fitting we solve for material parameters other than *B* and *α*.

The best-fit *G*-*γ* curves, along with the corresponding experimental data, are shown in Fig. 3. The optimal material parameters and the associated relative errors are shown in Tabs. S3 and S4 in the Supporting Materials. It can be seen from Fig. 3 that for both the collagen and fibrin networks, the shear modulus increases with shear strain, again indicating a nonlinear strain-stiffening behavior of the networks. As already mentioned, this behavior is induced by fiber realignment along the direction of the maximum principal stretch, and is reinforced by strain stiffening of individual fibers.

**Figure 3:**
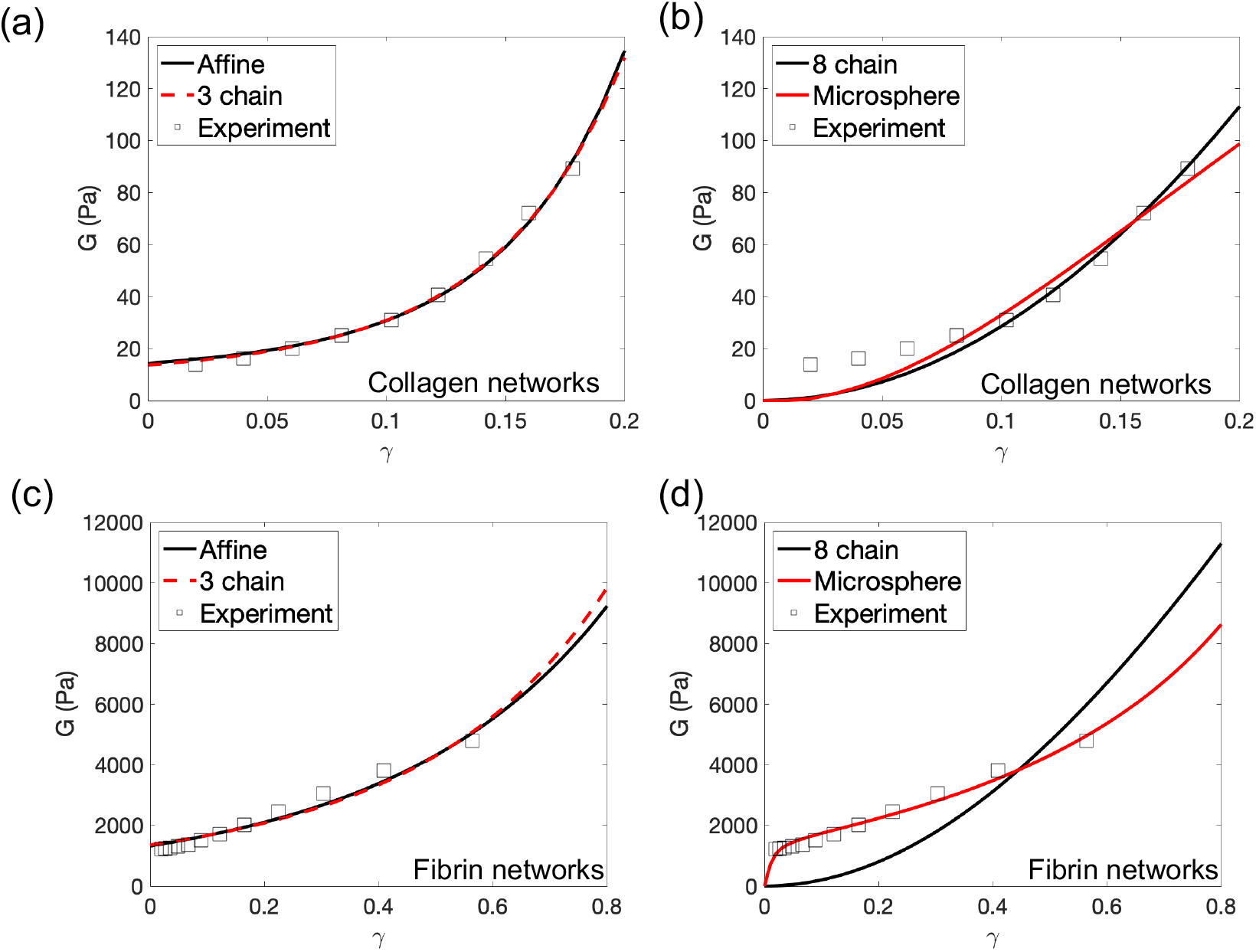
Nominal shear modulus *G* as a function of the amount of shear *γ*. Results predicted by the (a) affine and 3-chain models, as well as by the (b) 8-chain and micro-sphere models, are shown for collagen networks. The experimental data of Storm et al. is also displayed. The corresponding results are also shown for fibrin networks, including predictions of the (c) affine and 3-chain models, as well as of the (d) 8-chain and micro-sphere models. The corresponding experimental data of van Oosten et al. [32] is also shown.

From Fig. 3 we observe that both the affine and three-chain continuum models accurately describe the network shear response. In contrast, the eight-chain and micro-sphere models capture network response at large strains, but significantly underestimate the shear modulus at small strains. In particular, as the shear strain tends to zero, the shear modulus predicted by both models also tends to zero, indicating that the networks are mechanically unstable at small shear strains. These predictions significantly differ from the true network properties.

Upon closer inspection, we identify two reasons that explain why the eight-chain and micro-sphere models lead to unstable network response at small strains. First, in both models, under a prescribed network deformation all the fibers are stretched to the same degree described by a single, effective stretch (see (9) and (11)). (In contrast, in the affine and three-chain models, fibers along different directions are generally stretched to different extents, as would be expected in real situations.) Second, all fibers are assumed to be force free at zero strain. It can be shown (see the Appendix) that taken together, these two factors lead to a vanishing shear modulus at small strains.

In this context, we note that if all the fibers are taken to be initially pre-stretched, the eight-chain and micro-sphere models (as well as the affine and three-chain models) do give rise to stable network responses with positive shear moduli. However, it is known from more detailed, discrete fiber network simulations that prestress is *not* necessarily required for the stability of athermal networks [1, 3, 18, 44]. Therefore, the reliance of the eight-chain and micro-sphere models on prestress to achieve network stability is an unwanted feature, restricting their utility in describing the mechanical behavior of athermal networks.

Finally, we note that our results do *not* contradict earlier findings that the eight-chain and micro-sphere models can accurately describe the mechanical response of thermal networks like rubber. In those cases, chains within the network are always prestressed with an entropic, contractile tendency (unless the two ends of the chains overlap). As a result, the eight-chain and micro-sphere models are guaranteed to yield positive shear modulus and stable network behavior.

### 3.3. Combined tension and shear

In this subsection we study the performance of different models in characterizing network response under combined tension and shear. Specifically, we compare model predictions with the available experimental data of van Oosten et al. [32] for collagen and fibrin networks. In their experiments, cylindrical disk shaped network samples were sandwiched between the two parallel plates of a shear rheometer. These samples were axially compressed or stretched in steps by changing the gap between the two plates. At each level of axial stretch, the axial stress of the sample was measured after the fluid had re-distributed and the material had reached an equilibrium state. In addition, a small amount of shear strain was superimposed—by rotating one of the plates relative to the other. This was used to measure the shear modulus of the sample at that level of axial stretch. Thus, both the axial stress and the shear modulus were recorded as a function of axial stretch.

Given that the network samples were very thin, and that both the top and bottom surfaces of the samples were firmly attached to the rheometer plates, we assume that the lateral strain of the samples was zero. This assumption has also been used in previous studies to simulate the loading conditions in such experiments [18, 45]. Thus, we may write the deformation gradient as

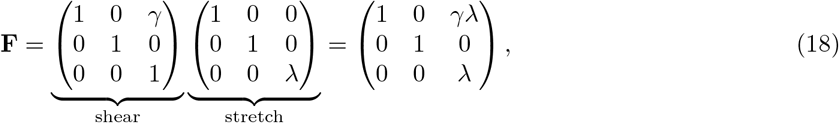

where *λ* is the axial stretch, and *γ* is the amount of superimposed shear, with *γ* = 2% in the experiment.

For each model, we first set *γ* = 0 in (18) and use (13) to compute the axial stress, *P*_*zz*_, as a function of *λ*. We then set *γ* = 2% and repeat the above procedure to compute the nominal shear modulus, *G* = *P*_*xz*_/*γ*, as a function of *λ*. Thereafter, we simultaneously fit the stress-stretch (*P*_*zz*_-*λ*) relation, as well as the modulus-stretch (*G*-*λ*) relation, to the corresponding experimental data for collagen and fibrin networks. The stress data and the modulus data have rather different magnitudes. To avoid bias in data fitting, in the cost function we use different weights for the two data sets. In particular, we choose weights that are inversely proportional to the squared magnitudes of the two data sets.

Figure 4 presents the best-fit curves and the corresponding experimental data for collagen networks, with Tab. S5 showing the calibrated material parameters and the associated relative errors. Figure 5 and Tab. S6 display the corresponding results for fibrin networks.

**Figure 4:**
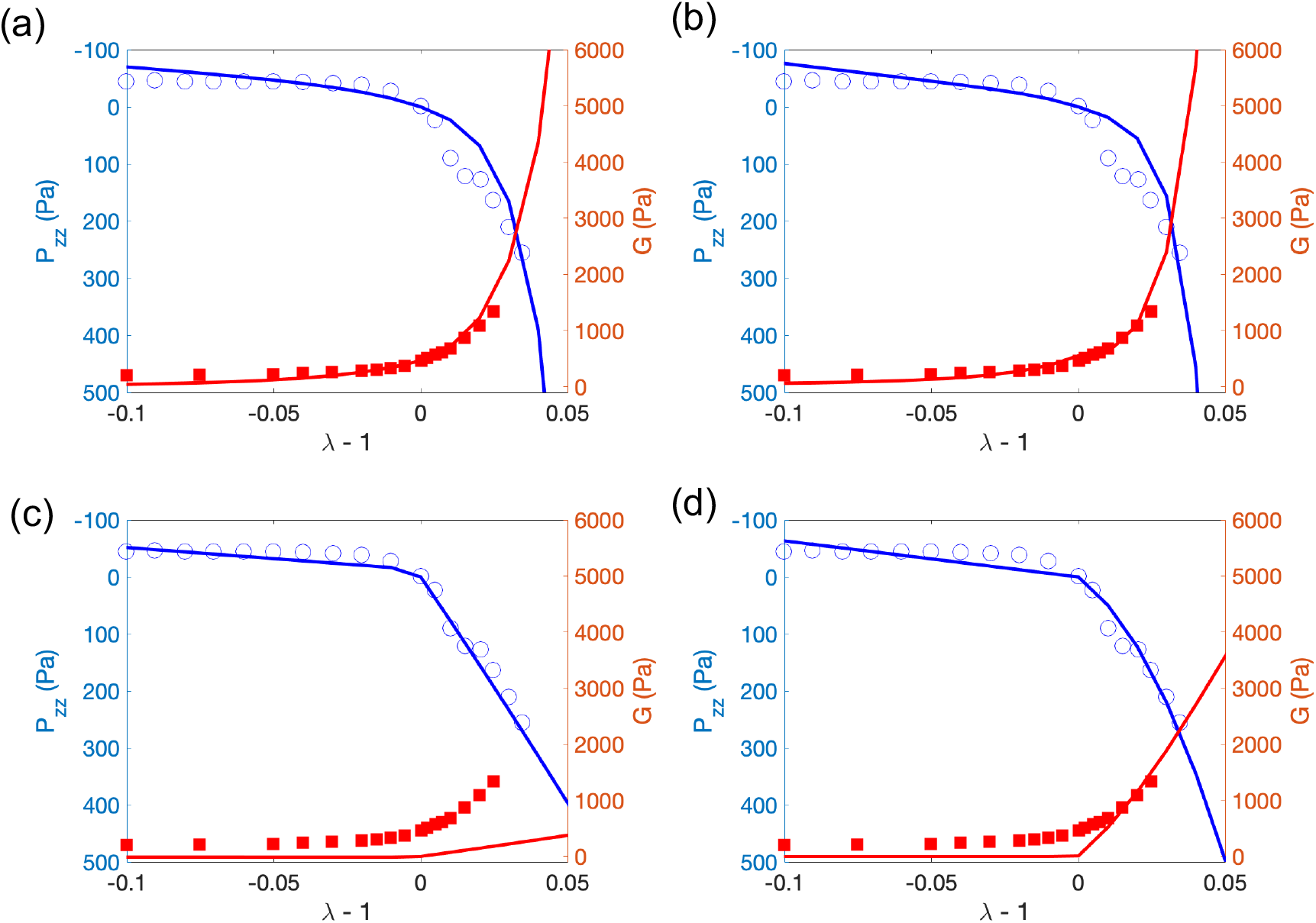
Axial stress *P*_*zz*_ and nominal shear modulus *G* of collagen networks as a function of axial strain (*ϵ* = *λ* − 1). Results predicted by the (a) affine, (b) 3-chain, (c) 8-chain, and (d) micro-sphere models, as well as the corresponding experimental results of van Oosten et al., are shown.

**Figure 5:**
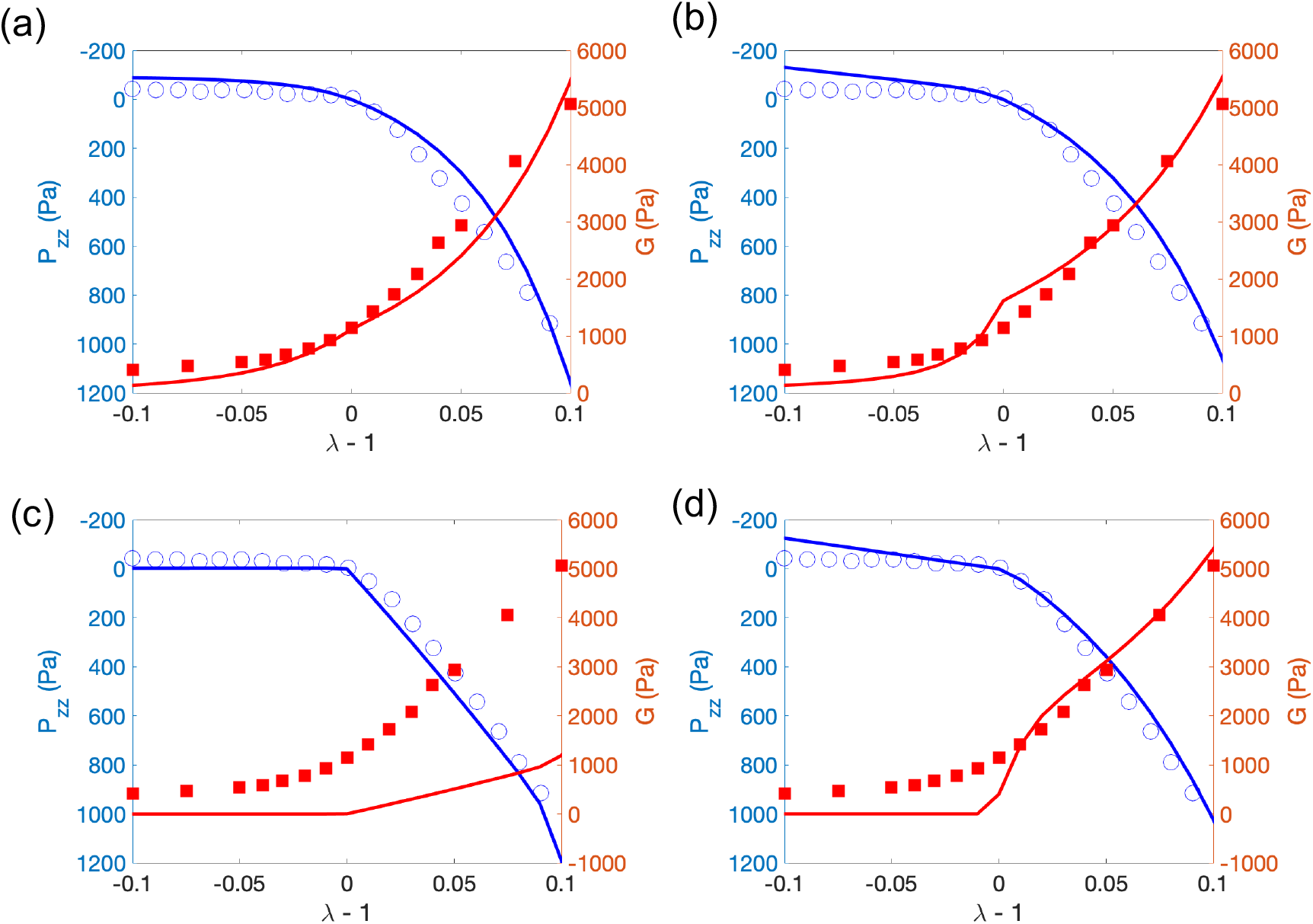
Axial stress *P*_*zz*_ and nominal shear modulus *G* of fibrin networks as a function of axial strain (*ϵ* = *λ* − 1). Results predicted by the (a) affine, (b) 3-chain, (c) 8-chain, and (d) micro-sphere models, as well as the corresponding experimental results of van Oosten et al., are shown.

It can be seen from Fig. 4 that the stress-stretch behavior of the collagen network displays strong tension-compression asymmetry, with network behavior being much stronger under tension than under compression. Correspondingly, the network shear modulus increases with increasing extension, but decreases with increasing compression. The stiffening of the network in tension can be attributed, once again, to fiber realignment along the loading direction and strain stiffening of fibers, whereas softening in compression is primarily due to buckling of fibers [18, 32].

Further, we observe from Fig. 4 that the affine and three-chain models can reasonably characterize both the axial and shear response of the network. On the other hand, the eight-chain and micro-sphere models capture the axial properties of the network, but not its shear properties. In particular, the eight-chain model significantly underestimates the networks shear modulus for all strains, and the micro-sphere model underestimates the shear modulus for small tensile and compressive strains. These discrepancies can again be attributed to the fact that these two models describe fiber deformations using a single effective stretch, which is insufficient to simultaneously capture the multi-axial response of the network. In particular, for the special case when the axial strain is zero (*λ* = 1), the deformation gradient in (18) reduces to simple shear in (17). In this case, the shear modulus predicted by the eight-chain and micro-sphere models will always be close to zero (as discussed in Section 3.2), thus failing to match the experimental data. We note that the predicted shear modulus is not exactly zero due to the finite amount of shear strain, i.e., 2%, applied to the network. As this shear strain tends to zero, the predicted shear modulus will tend to zero (as shown in the Appendix). We also make similar observations for the fibrin networks in Fig. 5.

In this context, we remark that the kinematics and mechanics of random athermal networks depend strongly on the detailed structure of the network, such as the network connectivity and cross link density [1, 2, 44]. In particular, deformation of fibers in random networks is known to be highly non-affine [8, 46, 47, 48], partially due to heterogeneities in network structure. These detailed structural features and complex fiber kinematics, are *not* captured by any of our continuum models, which all employ simplifying assumptions about network structure and fiber kinematics [3]. However, the affine and the three-chain models appear to have sufficient complexity that they are able to represent the macroscopic behavior of the networks with reasonable accuracy. Thus, they can be used as efficient, reduced-order tools to describe the macroscopic responses of athermal networks. In particular, these models can be easily implemented in finite-element packages to perform large-scale, structural simulations for athermal networks.

Similar to our findings, a recent study on modeling of fibrous tissues also suggests that, although the affine model fails to describe the underlying non-affine fiber kinematics, it can capture the macroscopic response of fibrous tissues using appropriate material parameters obtained from data fitting [49]. These material parameters, however, must be interpreted with care, because they are tuned to compensate for the model errors made in describing the network structure and kinematics. Consequently, their values (obtained from data fitting) can differ significantly from those that represent the true fiber properties.

Finally, we emphasize that our results do *not* imply that the affine and three-chain models are better than the eight-chain and micro-sphere models *per se*. In fact, as already mentioned, the eight-chain and microsphere models do a better job in characterizing the mechanical behavior of thermal networks like rubbers [19, 28]. Apparently, the relative performance of various network models depends on specific network types. Thus, we recommend that practitioners should systematically investigate the behavior of different models and select the ones that are best suited for the tasks at hand. Our study represents an example of one such effort.

## Conclusion

In this work, we developed continuum models to describe the compressible, hyperelastic behavior of isotropic athermal fibrous networks. For this purpose, we combined a single-fiber model that captures the asymmetric axial behavior of athermal fibers, with various network models—including the affine, three-chain, eight-chain and micro-sphere models—that assemble individual fiber properties into overall network response. Then, we systematically investigated the accuracy of these models by comparing model predictions with available experimental data for collagen and fibrin networks. Specifically, we tested how well each model captures the true network behavior under three loading conditions: uniaxial tension, simple shear, and combined tension and shear. We found that the affine and three-chain models can reasonably describe both the axial and shear response of the network. In contrast, the eight-chain and micro-sphere models accurately describe the axial behavior, but are inadequate in capturing the shear behavior. In particular, these models lead to zero shear modulus and thus to unstable network response at infinitesimal strains. We therefore recommended the use of the affine and three-chain models to describe the mechanical behaviors of athermal networks. These models can be implemented in finite-element codes to conduct large-scale, structural simulations for biopolymer networks like collagen and fibrin networks, serving as efficient tools to guide the design of fibrous scaffolds for cell engineering, and to understand the role of mechanics in pathologies.

## Acknowledgments

This material is based upon work supported in part by the U.S. National Science Foundation (NSF) under Grant Number CMMI-1548571, NSF 16-545, and DMR 17-20530. Dawei Song also acknowledges the support from the Center of Engineering and Mechanobiology (CEMB) Pilot Award. The authors declare no conflict of interest.

## Appendix

In this Appendix, we employ the eight-chain and micro-sphere models to compute the nominal shear modulus of athermal fibrous networks, which undergo simple shear deformations. In particular, we demonstrate that when all the fibers are initially force free, the shear moduli predicted by both models are zero at infinitesimal strains.

### A1. Eight-chain model

The first Piola-Kirchhoff stress predicted by the eight-chain model is given by (13), with *W*_1_ given by (10) and (9). Substituting the simple-shear deformation gradient (17) into (13), and using the definition of the nominal shear modulus, *G* = *P*_*xz*_/*γ*, we arrive at

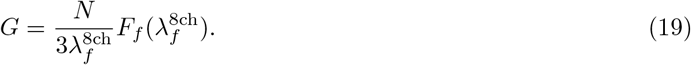

Here we recall that *N* is the number density of fibers, 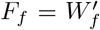 is a fiber’s axial force (see, e.g., (2)), and 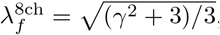, with *γ* being the amount of shear in (17). Then, it can be easily shown that

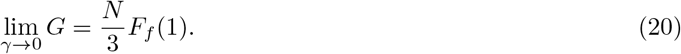

If all the fibers are initially force free (i.e., *F*_*f*_ (1) = 0), lim_*γ*→0_ *G* = 0, implying that the mechanical behavior of fibrous networks is initially unstable.

Note that if all the fibers are pre-stretched (i.e., *F*_*f*_ (1) > 0), lim_*γ*→0_ *G* > 0. This is indeed the case for thermal networks like rubbers, in which individual fibers are always pre-stretched with a contractile tendency (unless the two ends of a fiber coincide). On the other hand, if all the fibers are pre-compressed (i.e., *F*_*f*_ (1) < 0), lim_*γ*→0_ *G* < 0.

### A2. Micro-sphere model

The first Piola-Kirchhoff stress predicted by the micro-sphere model is given by (13), with *W*_1_ given by (12) and (11). Substituting the simple-shear deformation gradient (17) into (13), and using *G* = *P*_*xz*_/*γ*, we arrive at

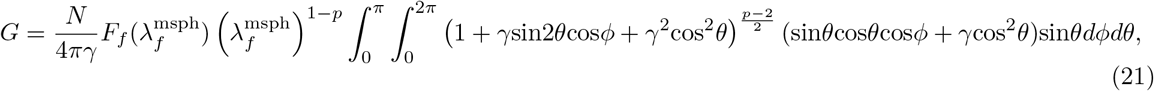

where we recall that *p* > 0 is a material parameter, and 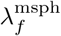 is given by

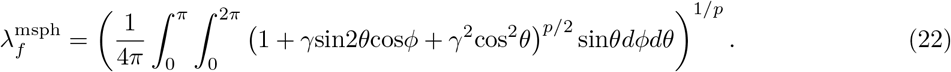

Note that for the special case *p* = 2, equations (21) and (22) can be evaluated analytically, and (21) can be shown to reduce to (19). However, for general values of *p*, (21) and (22) cannot be computed analytically and numerical integration is required.

Next, we compute the normalized modulus, *G*\* = (4*πG*)/(*Nκ*_0_), as a function of *γ*, and study how *G*\* behaves as *γ* → 0. (Recall that *κ*_0_ is a fiber’s stiffness at infinitesimal strains, see (1).) To do this, we need to know the axial responses of individual fibers. Since the value of *G*\* at infinitesimal strains is independent of the axial responses of fibers at large strains, we assume, for simplicity, that the fibers are linear elastic, that is *F*_*f*_ (*λ*_*f*_) = *κ*_0_(*λ*_*f*_ − 1). Other forms of *F*_*f*_ can also be used, but they will not affect the value of *G*\* as *γ* 0. We observe from Fig. 6 that lim_*γ*→0_ *G*\* = 0 for all values of *p*, again implying that the networks are initially unstable.

**Figure 6:**
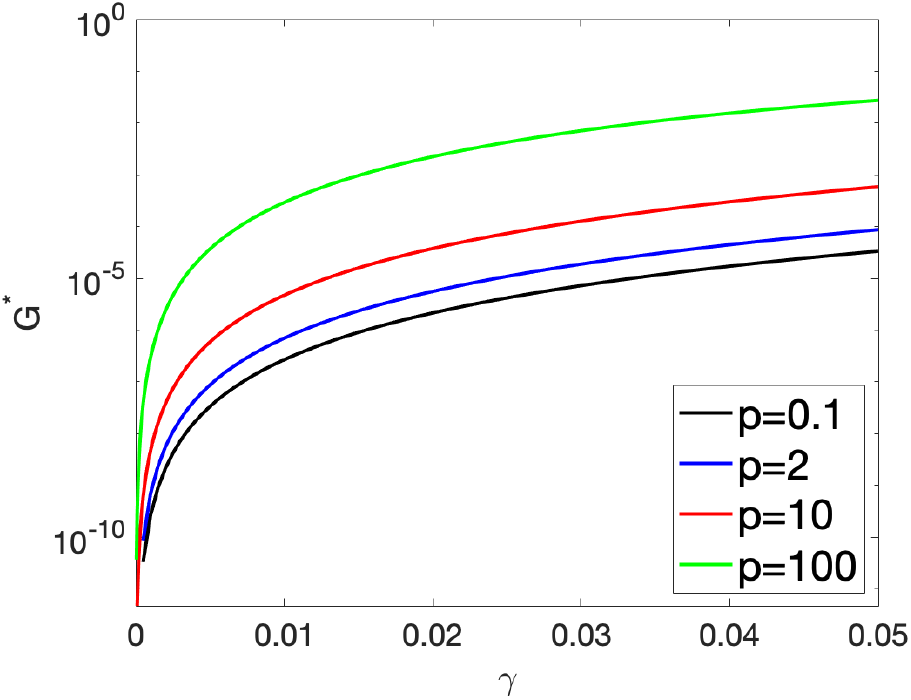
The normalized, nominal shear modulus *G*\* predicted by the micro-sphere model. Note that *G*\* = (4*πG*)/(*Nκ*_0_), with *G* given by equation (21). Regardless of the value of *p, G*\*→ 0 as *γ* →0, indicating that the fibrous networks are initially unstable.

