## Supplementary tables for "A study of hyperelastic continuum models for isotropic athermal fibrous networks"

| | **Affine** | **3-chain** | **8-chain** | **Micro-sphere** |
|---|---|---|---|---|
| $K$ | $30.15$ | $12.01$ | $298.12$ | $29.91$ |
| $d_0$ | $1e-2$ | $1e-2$ | $1e-2$ | $1e-2$ |
| $d_s$ | $0.197$ | $0.207$ | $853.78$ | $20.76$ |
| $B$ | $2.05e3$ | $5.13e3$ | $4.36e4$ | $152.06$ |
| $\alpha$ | $272.37$ | $267.34$ | $3.16e3$ | $279.08$ |
| $\varepsilon_s$ | $4.15e-6$ | $4.53e-6$ | $0.50$ | $0.25$ |
| $p$ | NA | NA | NA | $16.32$ |
| $e(c^*)$ | $8.66\%$ | $8.94\%$ | $13.26\%$ | $1.98\%$ |

Table S1. Material parameters obtained by fitting different models to the uniaxial-tension data of Roeder et al. [1] for collagen networks, along with the relative errors of data fitting.

| | **Affine** | **3-chain** | **8-chain** | **Micro-sphere** |
|---|---|---|---|---|
| $K$ | $183.0$ | $108.1$ | $327.1$ | $26.1$ |
| $d_0$ | $1e-2$ | $1e-2$ | $1e-2$ | $1e-2$ |
| $d_s$ | $0.599$ | $0.698$ | $0.450$ | $0.743$ |
| $B$ | $66.85$ | $33.24$ | $63.62$ | $108.65$ |
| $\alpha$ | $66.38$ | $13.44$ | $66.25$ | $58.67$ |
| $\varepsilon_s$ | $0.506$ | $0.514$ | $0.323$ | $0.248$ |
| $p$ | NA | NA | NA | $84.70$ |
| $e(c^*)$ | $3.22\%$ | $2.99\%$ | $5.32\%$ | $3.74\%$ |

Table S2. Material parameters obtained by fitting different models to the uniaxial-tension data of Purohit et al. [2] for fibrin networks, along with the relative errors of data fitting.

| | **Affine** | **3-chain** | **8-chain** | **Micro-sphere** |
|---|---|---|---|---|
| $K$ | $217.36$ | $81.94$ | $5.14e4$ | $1.40e3$ |
| $d_0$ | $2.14e3$ | $1.31e3$ | NA | $103.41$ |
| $d_s$ | $0.021$ | $0.025$ | $669.45$ | $103.41$ |
| $B$ | NA | NA | NA | NA |
| $\alpha$ | NA | NA | NA | NA |
| $\varepsilon_s$ | $1.20e-7$ | $4.18e-8$ | $0.73$ | $0.49$ |
| $p$ | NA | NA | NA | $32.33$ |
| $e(c^*)$ | $4.33\%$ | $3.66\%$ | $14.59\%$ | $14.80\%$ |

Table S3. Material parameters obtained by fitting different models to the simple-shear data of Storm et al. [3] for collagen networks, along with the relative errors of data fitting.

| | **Affine** | **3-chain** | **8-chain** | **Micro-sphere** |
|---|---|---|---|---|
| $K$ | $4.01e4$ | $1.63e4$ | $3.65e5$ | $6.80e3$ |
| $d_0$ | $9.88e-16$ | $1.37e-16$ | NA | $0.808$ |
| $d_s$ | $0.192$ | $0.192$ | $1.713$ | $0.222$ |
| $B$ | NA | NA | NA | NA |
| $\alpha$ | NA | NA | NA | NA |
| $\varepsilon_s$ | $6.47e-8$ | $7.87e-11$ | $0.052$ | $0.0083$ |
| $p$ | NA | NA | NA | $1.071e3$ |
| $e(c^*)$ | $8.76\%$ | $9.81\%$ | $52.03\%$ | $6.48\%$ |

Table S4. Material parameters obtained by fitting different models to the simple-shear data of van Oosten et al. [4] for fibrin networks, along with the relative errors of data fitting.

| | **Affine** | **3-chain** | **8-chain** | **Micro-sphere** |
|---|---|---|---|---|
| $K$ | $7e3$ | $3.33e3$ | $6.45e4$ | $3.18e4$ |
| $d_0$ | $0.015$ | $0.013$ | $5.90e-4$ | $2.64e-14$ |
| $d_s$ | $0.0097$ | $0.0089$ | $717.06$ | $1.14$ |
| $B$ | $397.05$ | $612.04$ | $398.05$ | $633.11$ |
| $\alpha$ | $0.0017$ | $3.23e-5$ | $0.0032$ | $2.94e-5$ |
| $\varepsilon_s$ | $0.001$ | $0.0065$ | $0.53$ | $0.47$ |
| $p$ | NA | NA | NA | $59.70$ |
| $e(c^*)$ | $26.79\%$ | $30.19\%$ | $64.90\%$ | $35.02\%$ |

Table S5. Material parameters obtained by fitting different models to the combined tension and shear data of van Oosten et al. [4] for collagen networks, along with the relative errors of data fitting.

| | **Affine** | **3-chain** | **8-chain** | **Micro-sphere** |
|---|---|---|---|---|
| $K$ | $1.72e4$ | $9.79e3$ | $8.69e4$ | $5.67e3$ |
| $d_0$ | $0.0176$ | $0.0097$ | $1.03e-4$ | $3.02e-15$ |
| $d_s$ | $0.0374$ | $0.0535$ | $0.0053$ | $0.0772$ |
| $B$ | $23.37$ | $992.09$ | $9.75e-18$ | $1.25e3$ |
| $\alpha$ | $6.28e-4$ | $4.44e-6$ | $248.28$ | $0.0012$ |
| $\varepsilon_s$ | $2.92e-14$ | $2.76e-14$ | $0.0289$ | $0.0086$ |
| $p$ | NA | NA | NA | $751.02$ |
| $e(c^*)$ | $16.40\%$ | $14.61\%$ | $59.26\%$ | $19.22\%$ |

Table S6. Material parameters obtained by fitting different models to the combined tension and shear data of van Oosten et al. [4] for fibrin networks, along with the relative errors of data fitting.


1. B. A. Roeder, K. Kokini, J. E. Sturgis, J. P. Robinson, and S. L. Voytik-Harbin, J Biomech Eng **124,** 214 (2002).
2. P. K. Purohit, R. I. Litvinov, A. E. Brown, D. E. Discher, and J. W. Weisel, Acta Biomater **7,** 2374 (2011).
3. C. Storm, J. J. Pastore, F. C. MacKintosh, T. C. Lubensky, and P. A. Janmey, Nature **435,** 191 (2005).
4. A. S. van Oosten, M. Vahabi, A. J. Licup, A. Sharma, P. A. Galie, F. C. MacKintosh, and P. A. Janmey, Sci Rep **6,** 19270 (2016).
